# Analysis of chemical and mechanical behaviors in living cells by continuum mechanics-based FRAP

**DOI:** 10.1101/2022.04.16.488540

**Authors:** Takumi Saito, Daiki Matsunaga, Shinji Deguchi

## Abstract

Fluorescence recovery after photobleaching (FRAP) is a common technique to analyze the turnover of molecules in living cells. Numerous physicochemical models have been developed to quantitatively evaluate the rate of turnover driven by chemical reaction and diffusion that occurs in a few seconds to minutes. On the other hand, they have limitations in interpreting long-term FRAP responses where intracellular active movement inevitably provides target molecular architectures with additional effects other than chemical reaction and diffusion, namely directed transport and structural deformation. To overcome the limitations, we develop a continuum mechanics-based model that allows for decoupling FRAP response into the intrinsic turnover rate and subcellular mechanical characteristics such as displacement vector and strain tensor. Our approach was validated using fluorescently-labeled beta-actin in an actomyosin-mediated contractile apparatus called stress fibers, revealing spatially distinct patterns of the multi-physicochemical events, in which the turnover rate of beta-actin was significantly higher at the center of the cell. We also found that the turnover rate is negatively correlated with the strain rate along stress fibers but, interestingly, not with the absolute strain magnitude. Moreover, stress fibers are subjected to centripetal flow as well as both contractile and tensile strains along them. Taken together, this novel framework for long-term FRAP analysis allows for unveiling the contribution of overlooked microscopic mechanics to molecular turnover in living cells.

## 1 Introduction

Fluorescence recovery after photobleaching (FRAP) is a widely used technique to evaluate molecular dynamics in living cells (1, 2). Time recovery of fluorescence intensity of target molecules within a bleached region is often fitted by a single exponential function to determine the characteristic time (3). The characteristic time estimates the involvement of individual physicochemical factors in the FRAP response. In chemical reaction-driven molecular turnover, for example, the fluorescence intensity is recovered without exhibiting diffusive gradients in space. Reaction rate equation-based models have been used in these cases to determine the kinetics (4, 5), in which the time constant in the single exponential function as a characteristic time gives the dissociation rate (4, 6). In diffusion-driven case where the Brownian motion-based pure-diffusion transports molecules, fluorescence distribution within the bleached region exhibits spatial gradients, meaning that the characteristic time depends on the size of the bleached region (7). The diffusion coefficient is then pragmatically estimated by considering the size of the bleached region and the time constant (8) or more accurately by analyzing the spatiotemporal fluorescence profile with a diffusion equation instead of single exponential functions (9–12). For more complicated cases with anomalous diffusion, fractional diffusion equation (13), alternative point FRAP (14), computational framework (15), and Fourier transform-FRAP with patterned photobleaching (16) have been developed to evaluate the effect of intracellular molecular crowding. Moreover, in a mixed case where diffusive molecules associate and dissociate with their reaction partners, mathematical models considering reaction-diffusion equations (4, 17, 18), FRAP combined with fluorescence correlation spectroscopy (19), and FRAP combined with genetic manipulation (7) have been developed to separately determine the diffusion coefficient and binding kinetics.

While many approaches have thus contributed to analyzing complex FRAP responses, long-term responses remain to be characterized. Specifically, the time course of intensity recovery inevitably displays an abnormal curve distinct from simple exponential functions because of the presence of intracellular directed movements. To analyze such abnormal curves, double exponential functions are instead used for determining two independent characteristic times (20, 21). However, the interpretation of the two characteristic times in terms of the actual physicochemical properties such as diffusion, directed movements, structural deformation, and chemical reaction is difficult. Regarding the involvement of directed movements in FRAP response, advection-reaction-diffusion models were developed to analyze one-dimensional movements of motor proteins in relatively short-term recovery curves of < 3 min (22, 23). In addition, advection-reaction models were developed to analyze two-dimensional directed movements of actin molecules observed in long-term FRAP experiments of > 10 min where fast diffusion process is ignorable (24). However, these models assumed constant velocities of the movement, and therefore more general nonlinearly and temporarily changing movements were beyond the scope of the analysis. Besides, intracellular structures are subjected to mechanical deformations or distortions, which are also not accessible with existing models. Consequently, it remains unclear how these unprobed effects, namely intracellular directed movements and structural deformations, affect the turnover within cells, which would be critical to understanding mechanotransduction at the organelle level to which these effects are related.

Here we develop a continuum mechanics-based approach to simultaneously determine the molecular turnover and active movements that inevitably appear in long-term FRAP responses. Analyses for such long-term behaviors are indispensable particularly for analyzing the dynamics of actin molecules associated with stress fibers, a contractile apparatus consisting of actin filaments, non-muscle myosin II, and other cross-linking and actin-binding proteins, given its long lifetime that exceeds 10 min (24–27). Because of the absence of appropriate FRAP analysis models for long-term behaviors, actin dynamics has not accurately been decomposed to evaluate the individual contribution of the two factors. Specifically, we obtain strain tensors of and turnover rate in moving photobleached areas, providing a basic framework to evaluate the turnover of molecules undergoing spatial movements and structural deformations.

## 2 Material and method

### 2.1 Cell culture, plasmids, and transfection

Rat aortic smooth muscle cell lines (A75r, ATCC) were cultured with low-glucose (1.0 g/L) Dulbecco’s Modified Eagle Medium (Wako) containing 10% (v/v) heat-inactivated fetal bovine serum (SAFC Biosciences) and 1% penicillin-streptomycin (Wako) in a humidified 5% CO_2_ incubator at 37 °C. An expression plasmid encoding mClover2-tagged beta-actin was constructed by inserting a human beta-actin gene, which was digested with XhoI and BamHI restriction enzymes from the EYFP-actin vector (#6902-1, Clontech) into the mClover2-C1 vector (Addgene plasmid #54577, a gift from Michael Davidson). The plasmid was transfected to cells using Lipofectamine LTX and Plus Reagent (Thermo Fischer Science) according to the manufacturer’s instructs.

### 2.2 FRAP measurements

Cells were cultured on a glass-bottom dish and transfected with the plasmid for 24 h for FRAP experiments. FRAP experiments were performed by using a confocal laser scanning microscope (FV1000, Olympus) with a 60X oil immersion objective lens (NA = 1.42). Fluorescence images were acquired every 10 s for 20 min using a 488-nm wavelength laser. Pre-bleach images were acquired for 20 s, and then photobleaching was carried out onto stress fibers for 3 frames by using a 405-nm and 440-nm wavelength laser.

### 2.3 Image analysis

Images were analyzed by using ImageJ (NIH) and MATLAB (MathWorks). Images were cropped to a tetragon to include a labeled region in single stress fibers. For the cropping, we set a threshold of intensity that automatically recognizes unbleached regions in the cropped pre-bleach frames, and thus the vertices of the labeled region were obtained in every time frame. For FRAP analysis, an intensity profile of the labeled region surrounded by the vertex points was spatially averaged over the area of A_bleach_, i.e.,

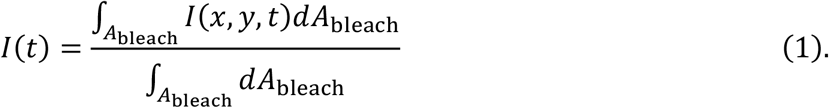

To describe the recovery curve, Eq. (1) was normalized by

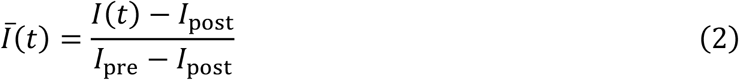

where *I*_pre_ and *I*_post_ represent the spatially averaged intensities in pre-bleach and bleached frames, respectively. The time course of normalized average intensity was fitted by using a least-square method with a single exponential function

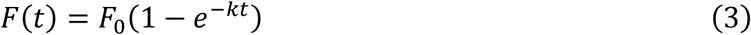

where *F*_0_ and *k* represent the mobile fraction and the turnover rate, respectively.

### 2.4 Continuum mechanics-based model of stress fibers

To evaluate mechanical properties of a labeled region by photobleaching on individual stress fibers, we first define position vectors corresponding to the vertex points forming the labeled region in cropped images (Fig. 1). The displacement vector between *t* = 0 (post-bleach) and *t* = *t*′ is described as

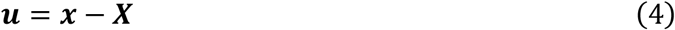

where ***X*** and ***x*** represent the position vectors at *t* = 0 and *t* = *t*′, respectively. The deformation of the labeled region is evaluated from the change in the displacement vectors to determine the displacement gradient *E*_*ij*_ (Green’s strain tensors) and *e*_*ij*_ (Almansi’s strain tensors), which are described by

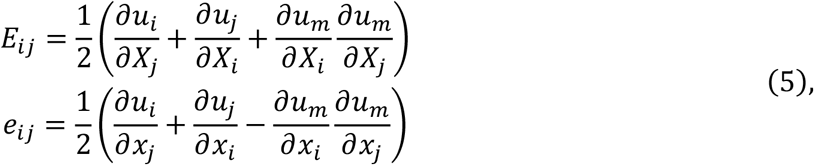

respectively. In the case of infinitesimal deformations, we can assume that *∂u*_*i*_*/∂X*_*j*_ ≃*∂u*_*i*_*/∂x*_*j*_, and the second order of the displacement gradient is negligible. The strain tensor is then reduced to

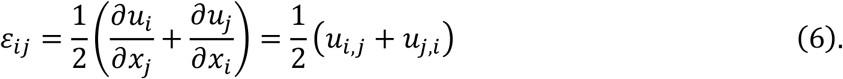

As the strain tensor is symmetric, coordinate-independent principal invariants are reduced to

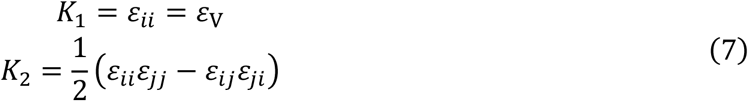

where *ε*_V_ is the volumetric strain. Substituting ***υ*** = *∂****u****/∂t* of the displacement rate for ***u*** of the displacement describes the strain rate tensor as

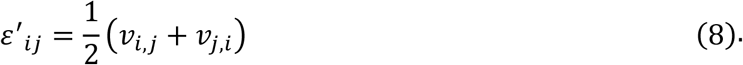

In the same manner, 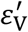 of the volumetric strain rate is obtained.

To distinguish between the deformation parallel and perpendicular to stress fibers, given vectors and tensors are transformed according to

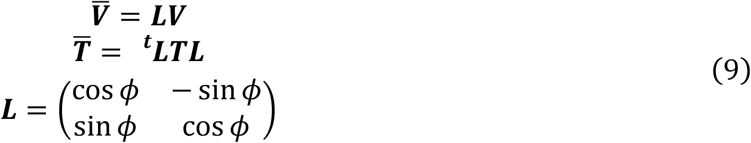

where ***V*** and ***T*** represent arbitrary vector and tensor, respectively, and ***L*** represents the transformation matrix determined by *ϕ* of the orientation of a single stress fiber within cropped images. The displacement and strain parallel and perpendicular to a stress fiber are then rewritten as

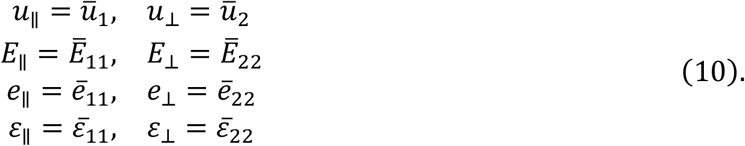

where the subscripts “‖” and “⊥” represent “parallel” and “perpendicular” to the stress fiber, respectively.

**Figure 1.**
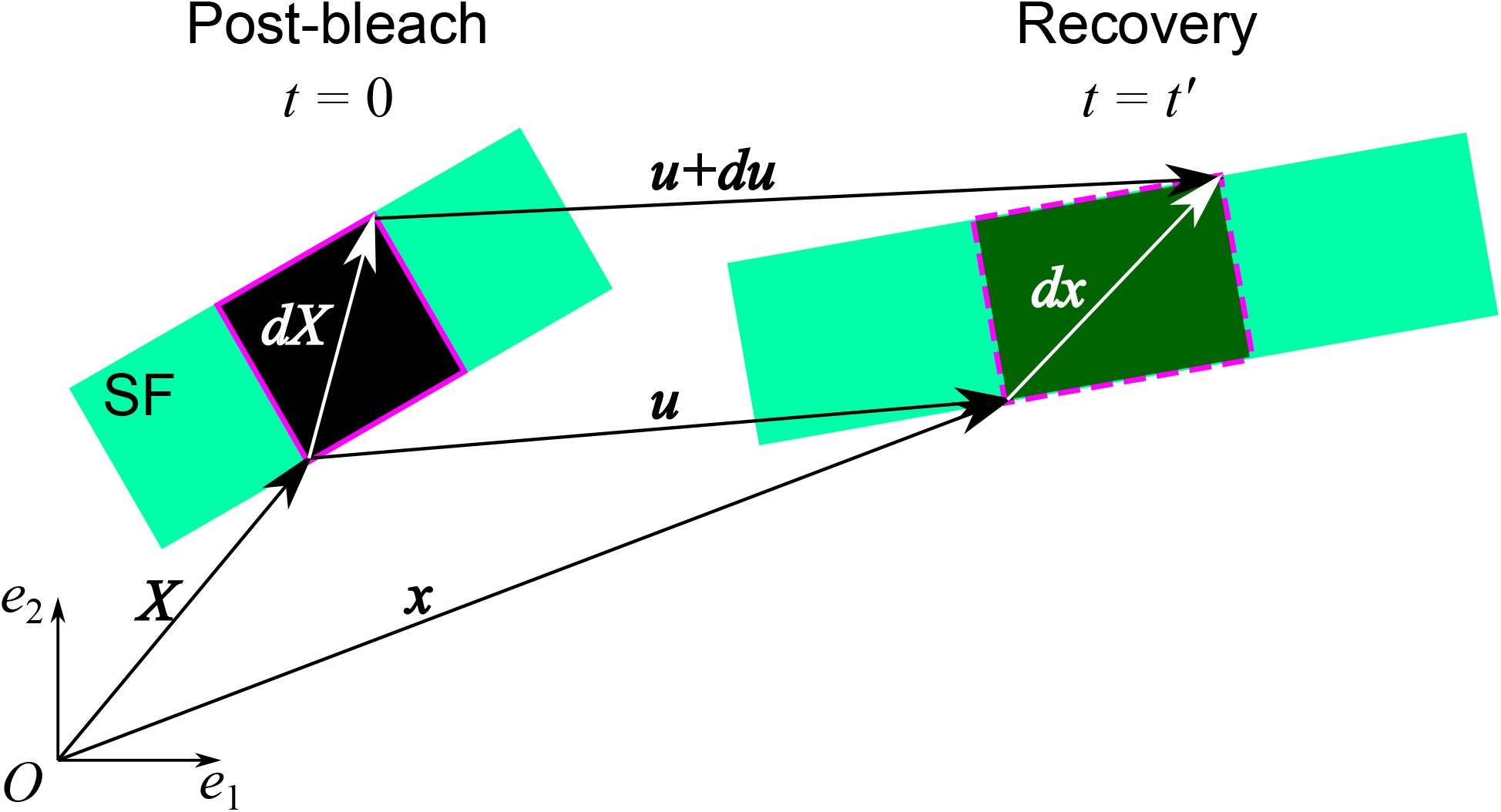
Schematic of the continuum mechanics-based FRAP model to analyze the chemical and mechanical properties within the region labeled by photobleaching. SF represents a part of a single stress fiber. See the text for details of the variables and coordinates.

### 2.7 Orientation analysis of stress fibers

Images were analyzed using ImageJ (NIH) and MATLAB (MathWorks). The orientation index between vectors is defined as

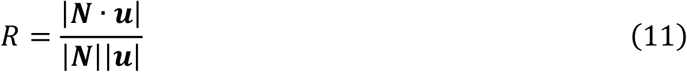

where *N* represents the orientation vector of a stress fiber. Here, *R* = 0 indicates that the displacement vector is perpendicularly oriented to the stress fiber, while *R* = 1 means the displacement vector is parallel with the stress fiber.

### 2.8 Statistical analysis

Unless otherwise stated, data are expressed as the mean ± standard deviation. Differences were calculated based on the Mann-Whitney U-test for variables with a non-Gaussian distribution, or the unpaired Student’s T-test for variables with a Gaussian distribution, with a significance level of *p <* 0.05 (*), *p <* 0.01 (**), or *p <* 0.001 (***).

## 3 Results

### 3.1 Continuum mechanics-based FRAP model determines both molecular turnover and mechanical properties in cells

Photobleaching was carried out onto mClover2-actin localized in stress fibers in A7r5 cells (Fig. 2A). The labeled region was automatically extracted based on an intensity threshold that distinguishes between bleached and unbleached regions in cropped images (Figs. 2A-C and Video S1). The actin molecular turnover recovered the intensity within the region. Meanwhile, the bleached square regions were subjected to spatial movements. To separately evaluate the molecular turnover and subcellular mechanical properties, first, the time evolution of the intensity within the tracked bleached region was normalized by Eq. (2) and then fitted by the least-square method with Eq. (3) (Fig. 2D), determining *k* of the turnover rate (Table 1). Second, the analysis of the spatial movement provides the displacement vector at given time points (Figs. 2B and 2C). The magnitude of the displacement, namely the norm of the displacement vector, was increased over time (Fig. 2E). To evaluate spontaneous deformations of the tracked region, the Green’s and Almansi’s strain tensors were obtained by using the rate change in the displacement vectors according to Eq. (5) (Figs. S1A and S1B). The error between these stain tensors, defined by ∣ (*E*_*ij*_ − *e*_*ij*_)/*E*_*ij*_ ∣, was within a range of 10^−2^ − 10^−1^ for both (*i, j*) = (1,1) and (*i, j*) = (1,2) and of 10^−1^ − 10for (*i, j*) = (2,2) (Fig. S1C). This result allows us to assume that the strain is infinitesimal and use the simplified form of the strain tensor shown in Eq. (6). Notably, *E*_*ij*_, *e*_*ij*_, and *ε*_*ij*_ are dependent on the coordinate system, meaning that individual components of these tensors in a certain region are unable to be compared to those in other labeled regions of stress fibers having different orientations in cells. To overcome this limitation, the volumetric strain corresponding to the first principal invariant of the strain tensor(6) was finally evaluated according to Eq. (7) (Fig. 2F).

**Figure 2.**
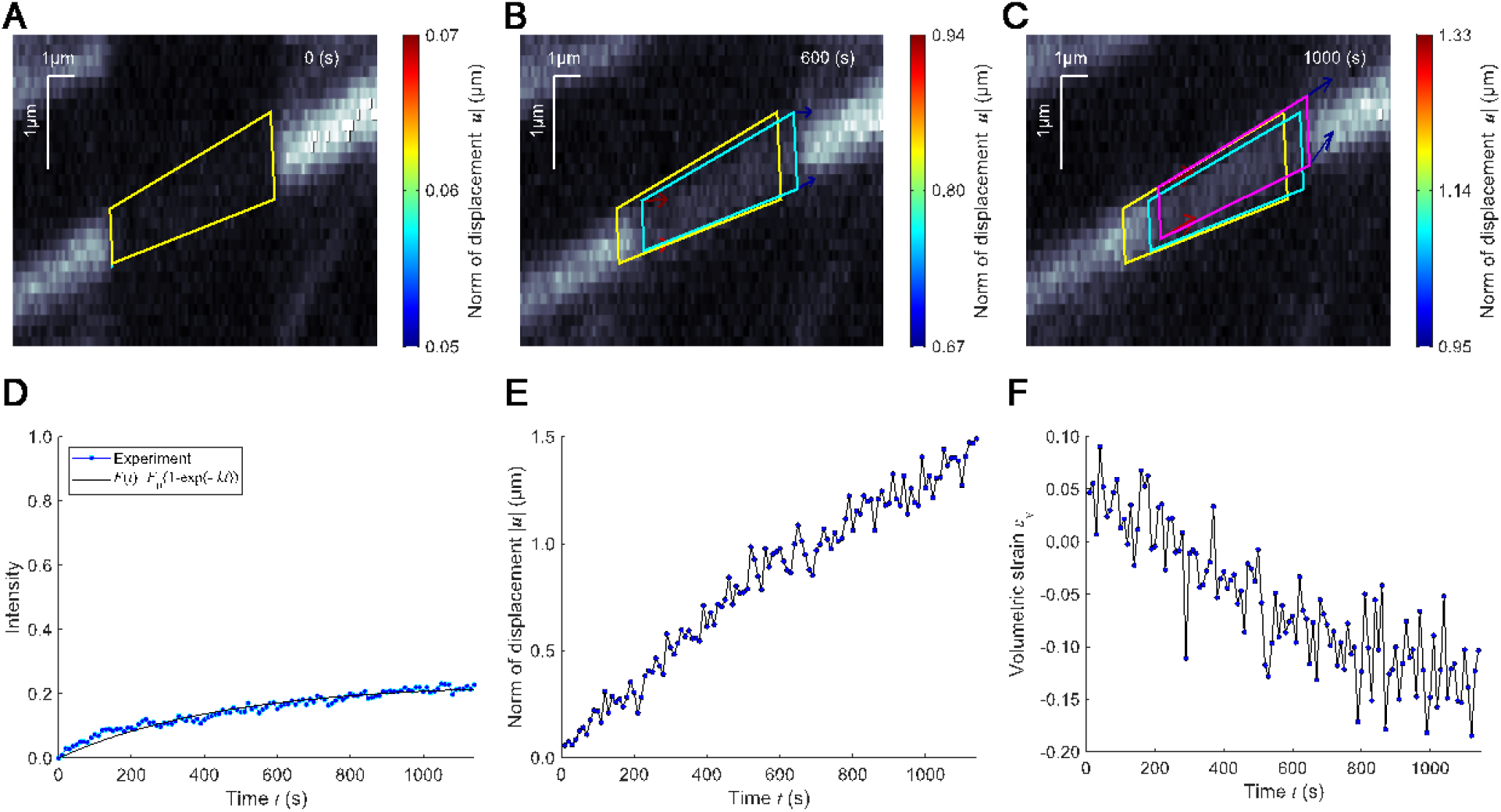
Determination of actin molecular turnover and mechanical properties (displacement and strain) of stress fibers in A7r5 cells. (A) The region of interest (ROI), automatically detected as shown in yellow, is labeled by photobleaching (*t* = 0 s). (B, C) The ROI, shown in cyan (*t* = 600 s) and magenta (*t* = 1000 s), is often spatially transported in a long-term observation (see Video S1). Arrows represent the tracked displacement vectors. (D) Time course of the fluorescence intensity within the ROI. The regression curve based on Eq. (3) determines the molecular turnover rate. (E) Time course of the norm of representative displacement vector. (F) Time course of the coordinate-independent volumetric strain.

**Table 1.** Turnover rate, displacement rate, and volumetric strain rate simultaneously determined.

| Turnover rate | Displacement rate | Volumetric strain rate |
| --- | --- | --- |
| $k$ (1/s) | $v$ ($\mu\text{m/s}$) | $\varepsilon'_v$ (1/s) |
| $(3.26 \pm 1.35) \times 10^{-3}$ | $(6.86 \pm 4.03) \times 10^{-4}$ | $(-1.29 \pm 1.41) \times 10^{-4}$ |

### 3.2 Mapping the distribution of mechanical behaviors in cells

To obtain the distribution of mechanical properties or more specifically intrinsic strain in cells, the displacement vector was calculated for multiple bleached regions on stress fibers (Figs. 3A-C and Video S2). Following this process, the volumetric strain, which is independent of the coordinates, was evaluated for these regions according to Eq. (6) and Eq. (7), providing the strain distribution in the cell (Figs. 3D-F and Video S3). The turnover rate of actin molecules was also determined by fitting Eq. (3) to fluorescence recovery curves of those regions (Fig. 2D). Our approach thus unveiled the mechanical properties as well as the molecular turnover along stress fibers in living cells (Table 1).

**Figure 3.**
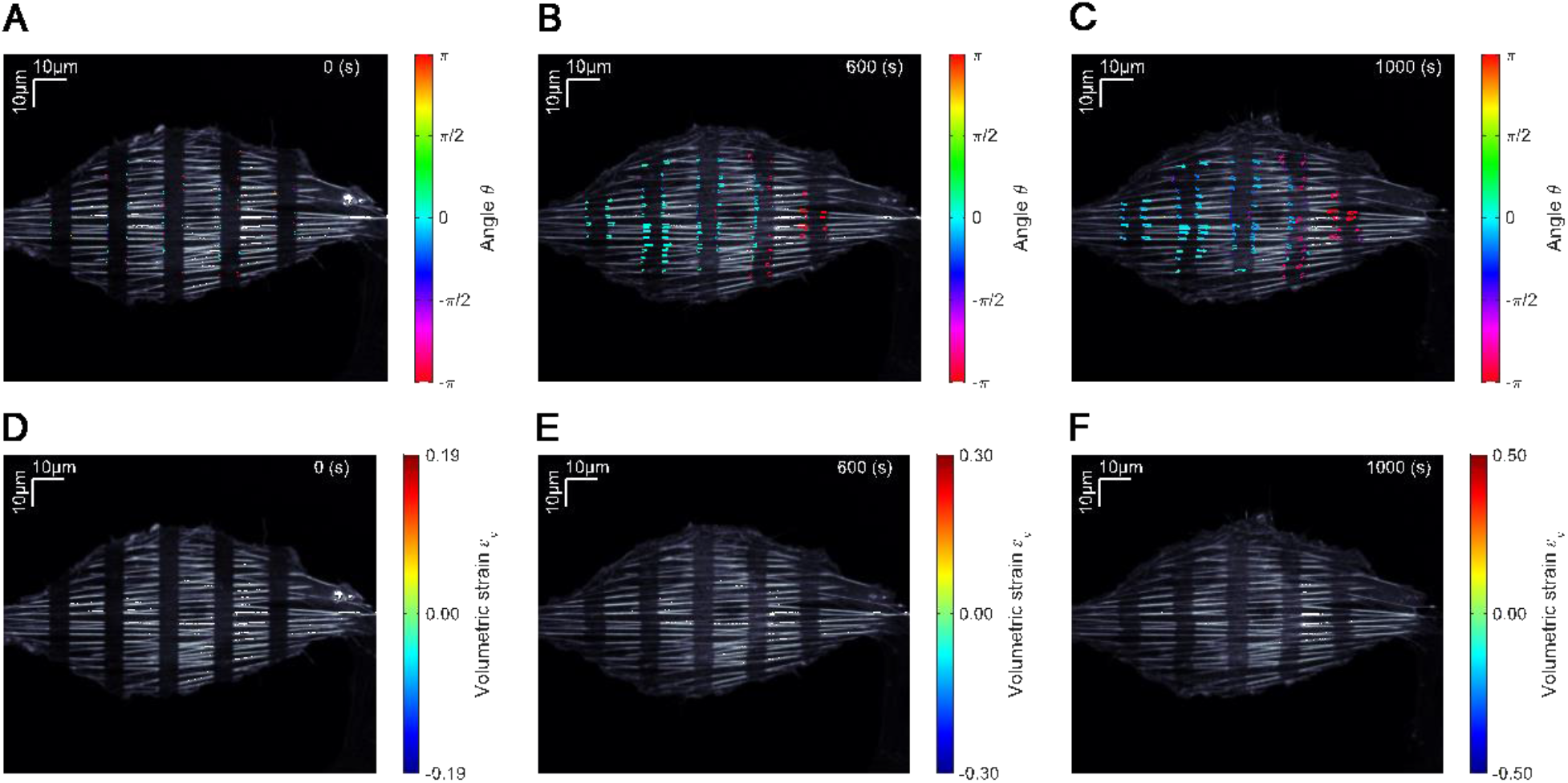
Multiple bleached regions on stress fibers in a single cell with the displacement and deformation distribution. Arrows and overlaid rectangular regions represent the displacement vectors and volumetric strain, respectively. (A-C) Time series of the displacement vectors. Color bar represents the angle of the displacement vectors (see Video S2). (D-E) Time series of the volumetric strain. Color bar represents the value of the volumetric strain (see Video S3).

### 3.3 Anisotropic mechanical properties distinct between parallel with and perpendicular to stress fibers

Stress fibers are subjected to non-muscle myosin II-dependent sustained contraction (28, 29). Probably because of this activity along the longitudinal direction of stress fibers, we found an anisotropic displacement that differs between the parallel and perpendicular directions (Figs. 3A-C). To evaluate the anisotropy, the orientation index *R*, which describes the correlation between displacement and orientation vectors of stress fibers, was calculated for multiple bleached regions according to Eq. (11), where *R* = 1 and *R* = 0 indicate the parallel and perpendicular correlation, respectively (Fig. 4A). The result showed that the displacement vector was predominantly parallel with the orientation of stress fibers as expected. The displacement vector was converted into the dominant coordinate system determined by the local orientation of single stress fibers in cropped images according to Eq. (9) and Eq. (10). The transformed vector parallel with stress fibers was more dominant than perpendicular one (Fig. 4B), thus consistent with the tendency of the orientation index.

**Figure 4.**
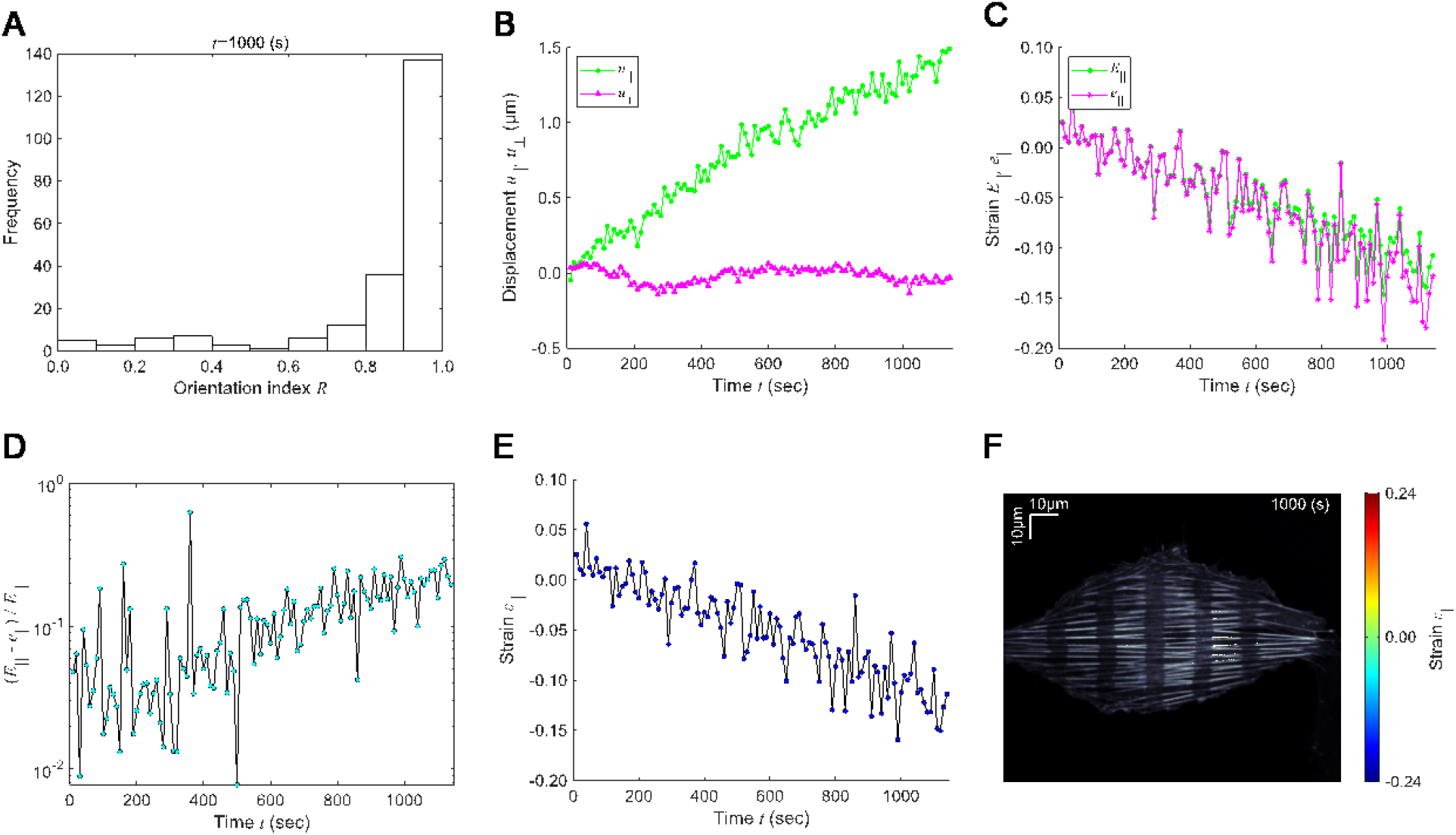
Long-term strain analysis of stress fibers. The displacement vectors and strain tensors are transformed to either parallel with or perpendicular to stress fibers. The subscripts “∥” and “⊥” represent parallel with and perpendicular to stress fibers, respectively. (A) The orientation index defined in Eq. (11) at *t* = 1000 s. (B) Time courses of the displacement vectors parallel with (green) and perpendicular to (magenta) stress fibers. (C) Time course of the parallel components of the Green’s (green) and Almansi’s (magenta) strain tensors. (D) Time course of the error defined as |(*E*_‖_ − *e*_‖_)/*E*_‖_|. (E) Time course of strain parallel with stress fibers. (F) The values of strain parallel with stress fibers in the reference frame are overlaid in the cell image.

Next, Green’s and Almansi’s strain tensors were likewise converted to conform to the dominant coordinate system, providing the deformation parallel with (Fig. 4C) and perpendicular to stress fibers (Fig. S2A). To assess whether the deformation of labeled regions is infinitesimal, we analyzed the error defined by either |(*E*_‖_ − *e*_‖_)/*E*_‖_| or |(*E*_⊥_ − *e*_⊥_)/*E*_⊥_|. The difference between the Green’s and Almansi’s strains parallel with stress fibers remained below the magnitude of their values (Fig. 4D), while the magnitude of the displacement was increased over time (Fig. 4B). In contrast, the difference between their perpendicular components was almost the same as the magnitude of strains (Fig. S2B) probably because the small width of stress fibers, i.e., the perpendicular component of transformed 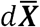 and 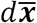, would include inevitable observation errors that could be comparable to the actual deformation. The strains along the width direction may therefore be affected particularly regarding the third term in Eq. (5), even though the displacement along the width of stress fibers remained within a range (Fig. 4B). Contrary to the behavior of the Green’s strain, the Almansi’s strain along the width of stress fibers was decreased with increasing time due to the opposite sign of the third term in Eq. (5) (Fig. S2A). On the other hand, by considering *u*_⊥_ of the small displacement compared to *u*_‖_ (Fig. 4B) in addition to the small difference between Green’s and Almansi’s strains parallel with stress fibers (Fig. 4C and 4D), we found that the deformation of labeled regions was infinitesimal to allow us to use 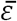 of the transformed strain tensor. The strains parallel with (Fig. 4E) and perpendicular to stress fibers (Fig. S3A) were then recalculated according to Eq. (9) and Eq. (10). We also obtained the ratio of the error between the strains parallel and perpendicular to stress fibers defined by |*ε*_⊥_/*ε*_∥_|, resulting in that the strain parallel with stress fibers was dominant (Fig. S3B), which agrees with the displacement vectors (Fig. 4A). The strain parallel with stress fibers was then overlaid onto the multiple regions to visualize the anisotropic mechanical properties of the cell (Fig. 4F and Video S4).

### 3.4 Actin molecular turnover increases at the cell center as opposed to mechanical behavior

Our results indicated that stress fibers are indeed subjected to the deformation that occurs predominantly along their longitudinal direction (Fig. 4A and 4B). However, it has remained an open question whether there is a cooperative or competitive effect between such mechanical behavior and actin molecular turnover on stress fibers. To decipher this complex process, we performed the FRAP experiments with the stripe bleach pattern that is approximately perpendicular to the dominant orientation of stress fibers in A7r5 cells (Fig 5A). FRAP responses within multiple bleached regions were analyzed at all the stripes from S1 to S5 to determine *k* of the turnover rate (Fig. 5B), *v*_‖_ = *u*_‖_/*t* of the velocity (Fig. 5C), and *ε*_‖_ of the strain (Fig. 5D). The turnover rate of actin in peripheral stripes of S1 and S5 was approximately two-fold slower than that of S3, increasing at the center of the cell (Fig. 5B). By contrast, the magnitude of velocity and strain along the stress fibers decreased at the center of the cell. Moreover, negative sign of the velocity on either side of the cell appeared while positive sign did on the other side, suggesting the presence of a centripetal flow (Fig. 5C). In a similar manner to the velocity, approximately one-half of cells tend to possess compressive strains whereas the others do tensile strains (Fig. 5D).

**Figure 5.**
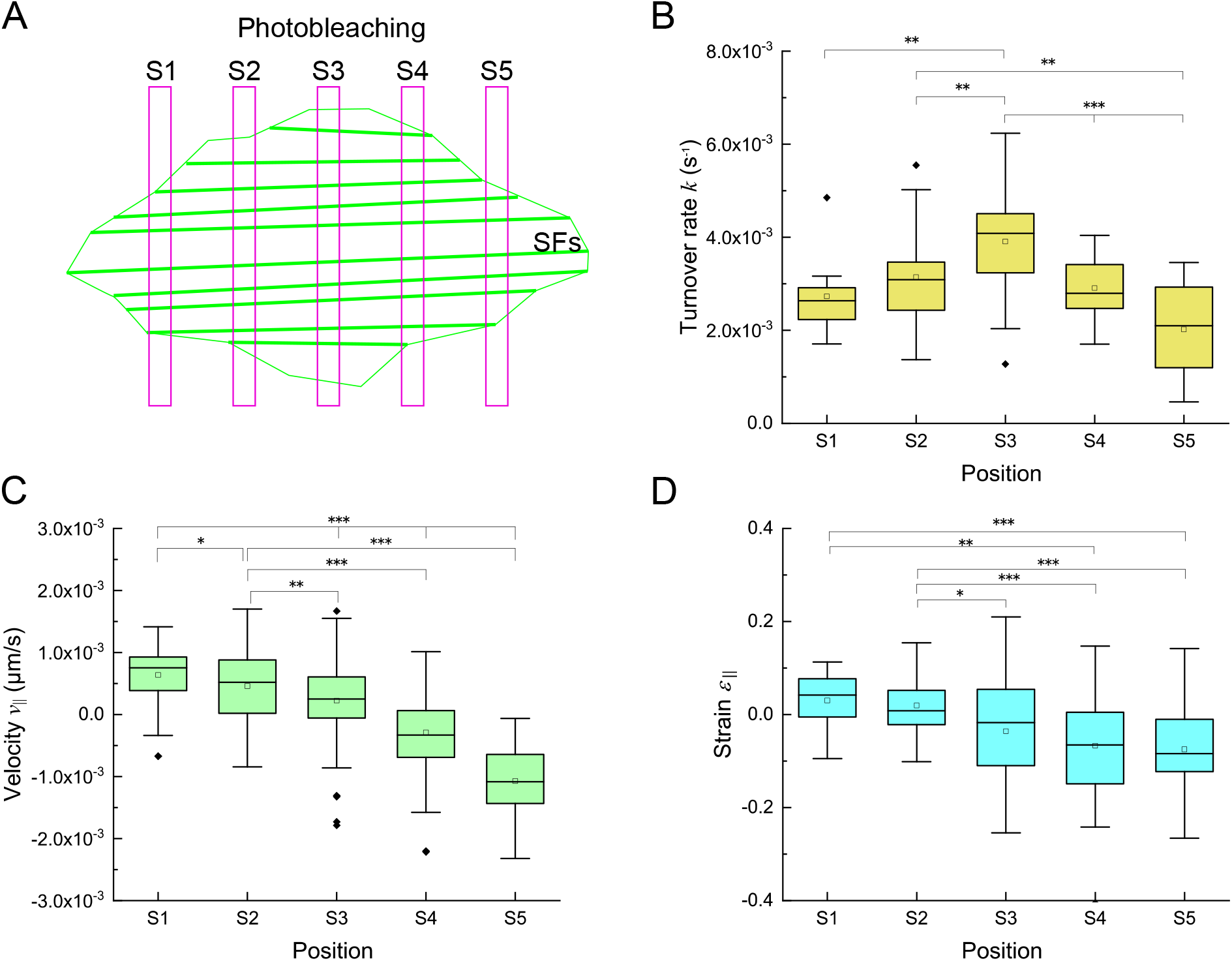
The chemical turnover and mechanical strain are high at the cell center and periphery, respectively. (A) Schematic of stripe pattern of photobleaching in a cell. (B-D) Actin turnover rate (B), *v*_‖_ = *u*_‖_/*t* of the displacement rate (C), and strain (D) along stress fibers are shown by mean ± SD analyzed at each stripe (*n* = 13, 34, 32, 32 and 15 for S1, S2, S3, S4 and S5, respectively). Asterisks represent a statistically significant difference among the stripes (*, *p <* 0.05; **, *p <* 0.01; ***, *p <* 0.001).

## 4 Discussion

Numerous FRAP methods have been proposed to evaluate the complex turnover in a normal diffusion-driven case (8–12), anomalous diffusion-driven case (13–16), chemical reaction-driven case (4, 5), and diffusion-reaction case (4, 7, 17). However, because of the current lack of appropriate FRAP methods, the “long-term” turnover driven by stress fiber-associated active dynamics that include both intracellular bulk-like flow and microscopic structural deformation remains highly elusive. The main difficulty is that fluorescence recovery curves exhibit abnormal curves unlike the conventional single exponential form due to spatial movements of labeled regions. To overcome this problem, double exponential functions are alternatively used to determine two independent characteristic times (20, 21). However, it is difficult to interpret such independent characteristic times in terms of the real physicochemical factors such as chemical reaction, spatial movements, and/or structural deformation that potentially all affect the long-term turnover. We here provided the continuum mechanics-based FRAP framework to reveal the multi-physicochemical contributions to the long-term turnover.

To demonstrate our approach, FRAP experiments were carried out for mClover2-tagged actin molecules on stress fibers in A7r5 cells. The material points forming the region of interest provide the position and displacement vectors at given time points (Fig. 1 and Fig. 2A-C). The rate change in displacement vector gives us the strain tensors, *E*_*ij*_, *e*_*ij*_, and *ε*_*ij*_. Notably, intracellular fibrous structures such as stress fibers exist with respective orientations in cells, meaning that the coordinate-dependent form is not adequate to extract vectors and tensors in multiple bleached regions. To circumvent this situation, the coordinate-independent volumetric strain, one of the principal invariants of the strain tensor, is determined (Fig. 2F). Meanwhile, the time evolution of intensity within the labeled region determines the molecular turnover rate (Fig. 2D). Taken together, our approach enables to evaluate comprehensive physicochemical properties, namely the chemical reaction-driven molecular turnover rate and the subcellular mechanical displacement and deformation.

Furthermore, we performed FRAP experiments with the stripe pattern to evaluate the distribution of mechanical and chemical properties on stress fibers in A7r5 cells (Fig. 5). By analyzing spatiotemporal profiles of multiple bleached regions, we made the following findings: (i) the actin turnover rate in peripheral regions of cells is two-fold faster than that at the center of the cell as suggested before regarding the faster turnover of actin cross-linking proteins in stress fibers (30), and (ii) both the velocity and the strain are decreased at the center of the cell in an opposite manner to the turnover rate (Fig. 5C and 5D).

It was an open question whether there is a cooperative or competitive interaction between the actin molecular turnover and such microscopic dynamics on stress fibers. To decipher this complex process, we analyzed the Pearson correlation coefficient between the actin molecular turnover rate and the magnitude of either the velocity of spatial displacement or the strain of microscopic structural deformation. We found a negative correlation *R*(*k*, |*v*_‖_|) = −0.34 with *p* = 0.18 × 10^−3^ (Fig. 6A), and no correlation *R*(*k*, |*ε*_‖_|) = 0.06 with *p* = 0.54 (Fig. 6B). Since the turnover rate in the chemical reaction-driven case is known to be involved in the off-rate (4, 6), the negative correlation of *R*(*k*, |*v*_‖_|) suggests that the dissociation of actin molecules with stress fibers is reduced by the mobility of stress fibers. Regarding the specific spatial distribution (Figs. 5B and 5C), it is likely that actin polymerization at the cell periphery may directly facilitate the spatial transport of stress fibers toward the cell center as is often observed in the retrograde flow (31, 32). Thus, turnover is rather suppressed to end up having locally stabilized stress fibers when they are under fast movement within cells. In contrast, no correlation between the turnover rate and the strain along the stress fibers suggests that the “microscopic deformation” is not a measure to evaluate the actin molecular turnover. Considering the microscopic structure of the contractile unit of stress fibers known as “non-muscle sarcomere” (27), actin filaments are basically only sliding to change the relative position with another stress fiber constituent non-muscle myosin II (28) in a manner similar to the actomyosin-mediated contraction in cytokinesis (33, 34). Thus, the net strain of individual actin filaments would not be necessarily comparable in magnitude to, or even be associated with, the microscopic strain observed at the whole cell level. Altogether, our findings made with the quantitative decoupling of the multiple-physicochemical phenomena suggest that stress fibers are transported toward the cell center where the turnover of actin molecules is accelerated, and stress fibers under such fast transportation are rather stabilized, while the macroscopic deformation or the absolute strain magnitude does not actually reflect the extent of the chemical reaction or turnover, thus giving novel insights into the complex cellular mechanism of how stress fiber-associated molecules respond to mechanical factors.

**Figure 6.**
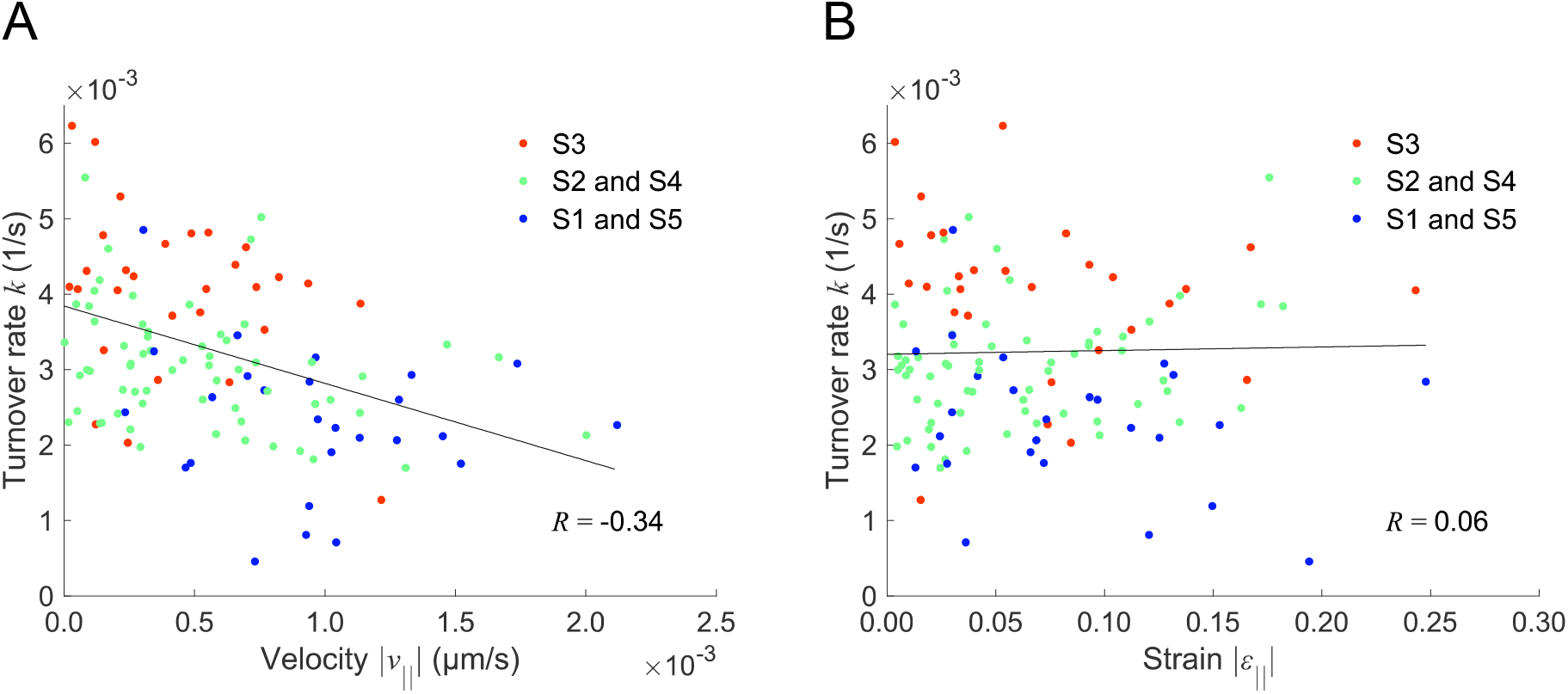
Correlation analysis represents the relationship between the molecular turnover and mechanical behaviors. The actin molecular turnover is negatively correlated with the magnitude of velocity parallel with stress fibers (*R*(*k*, |*v*_‖_|) = −0.34; *p* = 0.18 × 10^−3^) (A) but not with the magnitude of strain parallel with stress fibers (*R*(*k*, |*ε*_‖_|) = 0.06; *p* = 0.54) (B). Lines represent the regression fitting for variables of S1 and S5 (blue), S2 and S4 (green), and S5 (red).

## Acknowledgments

TS is supported by Japan Society for the Promotion of Science (JSPS). This study was partly supported by JSPS KAKENHI Grants (18H03518, 19K22967, and 20J10828).

## Supplementary Material

**Figure S1.**
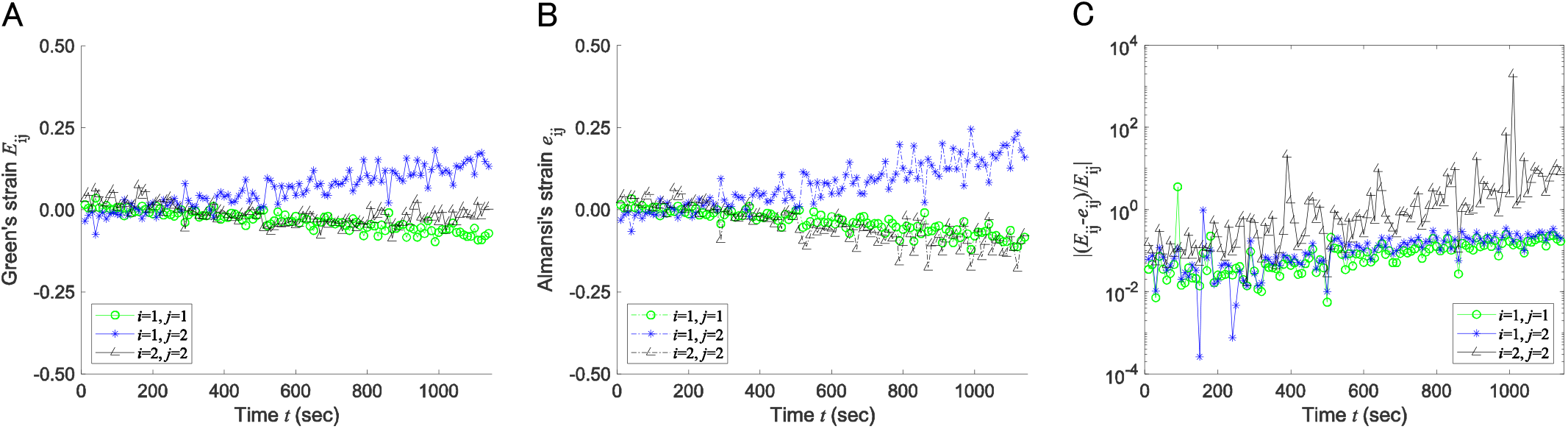
Green’s and Almansi’s strain tensors are determined by Eq. (5). (A) Time course of the components of Green’s strain tensor. (B) Time course of the components of Almansi’s strain tensor. (C) Time course of the error defined as ∣ (*E*_*ij*_ − *e*_*ij*_)/*E*_*ij*_ ∣.

**Figure S2.**
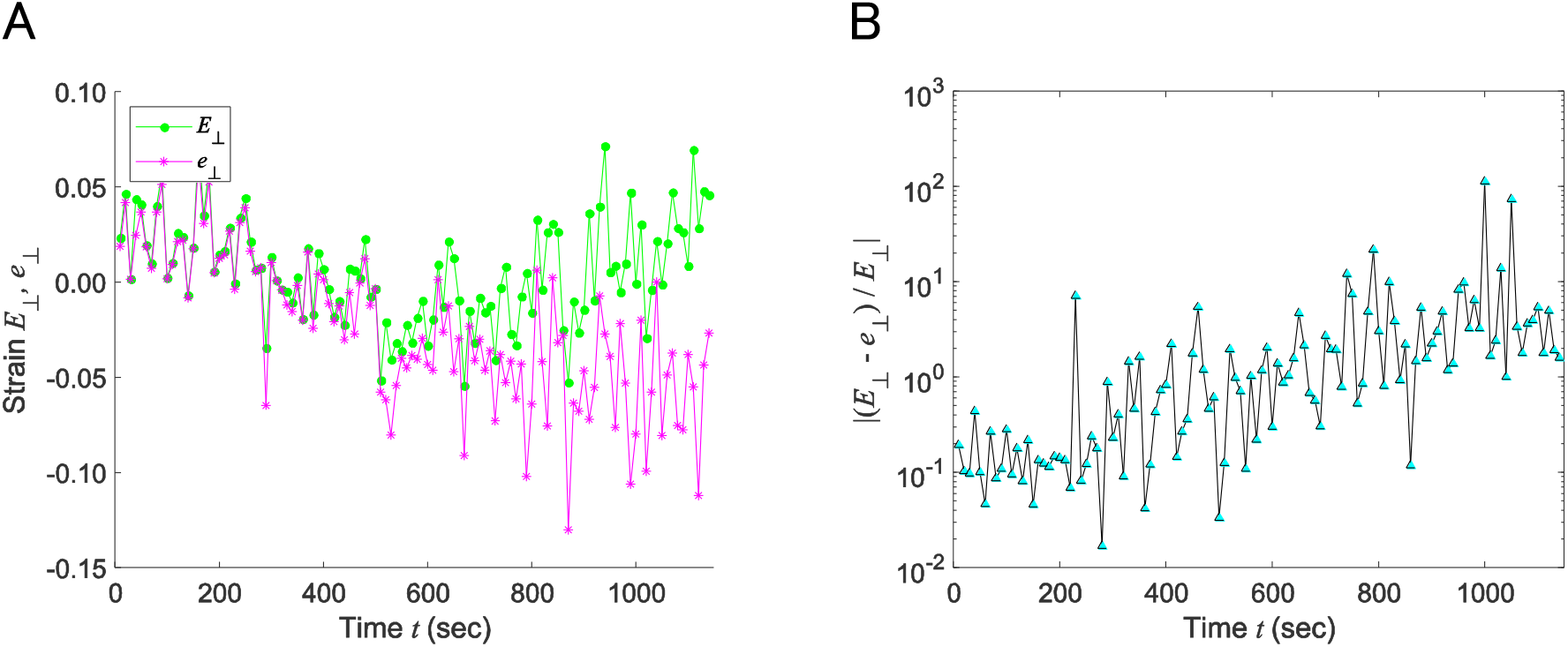
Green’s and Almansi’s strain tensors are transformed by Eq. (9). (A) Time courses of Green’s (green) and Almansi’s (magenta) strain perpendicular to stress fibers. (B) Time course of the error defined as |(*E*_⊥_ − *e*_⊥_)/*E*_⊥_|.

**Figure S3.**
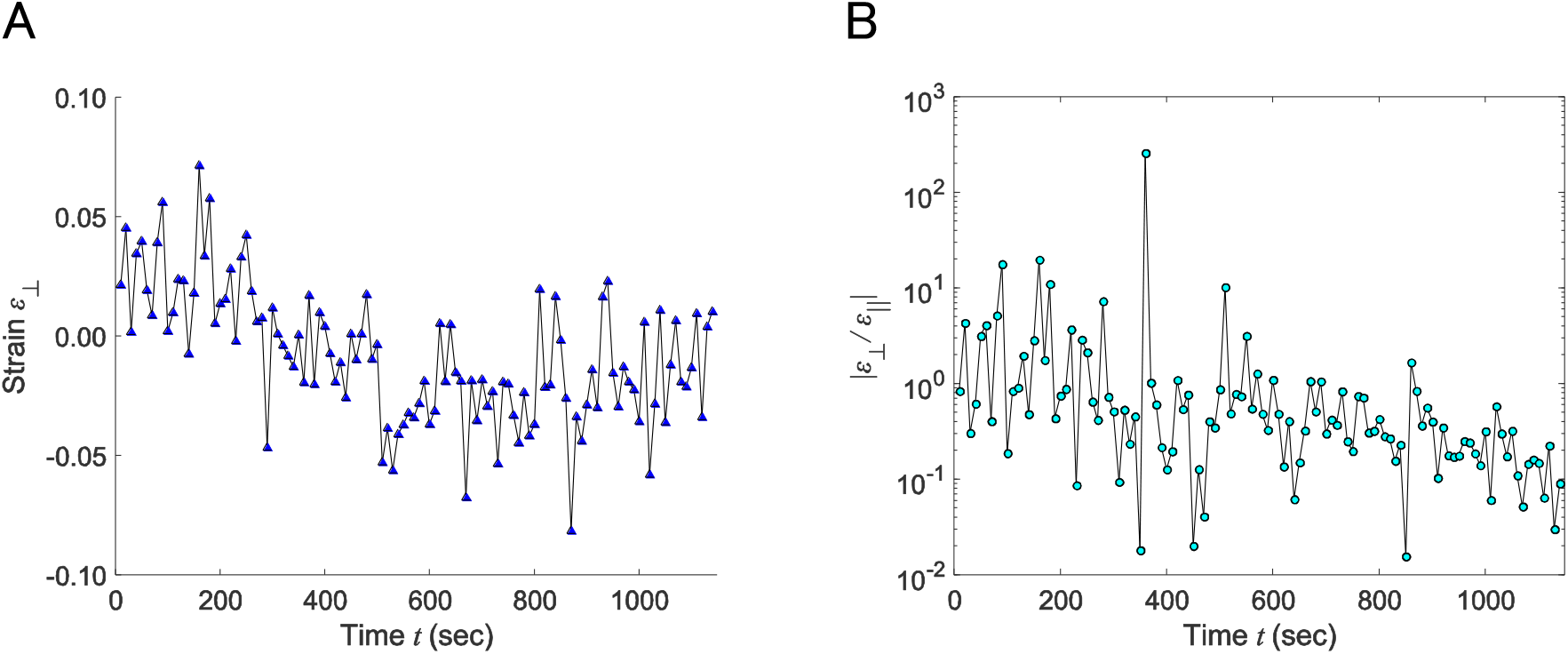
Strain perpendicular to stress fibers is ignorable compared to that along the axial direction. (A) Time course of strain perpendicular to stress fibers. (B) Time course of the ratio of perpendicular to parallel strain.

Video S1 FRAP images in a single stress fiber with a reference region.

Video S2 FRAP images with the displacement vectors analyzed at multiple bleached regions in a cell. Arrows represent the displacement vectors.

Video S3 FRAP images of the volumetric strain analyzed at multiple bleached regions in a cell.

Video S4 FRAP images of the strain along stress fibers obtained at multiple bleached regions in a cell.

